# Gametic specialization of centromeric histone paralogs in *Drosophila virilis*

**DOI:** 10.1101/530295

**Authors:** Lisa E. Kursel, Harmit S. Malik

**Affiliations:** Molecular and Cellular Biology Graduate Program, University of Washington, Seattle, USA; Division of Basic Sciences; Howard Hughes Medical Institute, Fred Hutchinson Cancer Research Center, Seattle, USA

## Abstract

In most eukaryotes, centromeric histone (CenH3) proteins mediate the highly conserved process of chromosome segregation as the foundational kinetochore assembly factor. However, in multicellular organisms, CenH3 proteins have to perform their essential functions in different chromatin environments. CenH3 proteins not only mediate mitosis and meiosis but also ensure epigenetic inheritance of centromere identity on sperm chromatin, which is highly compact and almost completely stripped of histones during spermiogenesis. We hypothesized that such disparate chromatin environments might impose different functional constraints on CenH3. If so, gene duplications could ameliorate the difficulty of encoding divergent and even potentially incompatible centromeric functions in the same gene. Here, we analyzed the cytological localization of two recently identified CenH3 paralogs, Cid1 and Cid5, in *D. virilis* using specific antibodies and epitope-tagged transgenic strains. We find that only ancestral Cid1 is present in somatic cells, whereas both Cid1 and Cid5 are expressed in testes and ovaries. However, Cid1 and Cid5 are alternately retained in male and female gametes; Cid1 is lost in male meiosis but retained throughout oogenesis, whereas Cid5 is lost during female meiosis but retained in mature sperm. Following fertilization, maternally deposited Cid1 rapidly replaces paternal Cid5 during the protamine-to-histone transition. Our studies reveal mutually exclusive gametic specialization of two divergent CenH3 paralogs. We suggest that centromeric histone duplication and divergence may allow essential genes involved in chromosome segregation to specialize and thereby resolve an intralocus conflict between maternal and paternal centromeric histone requirements in many animal species.

## Introduction

Chromosome segregation is an essential process that is highly conserved across eukaryotes. Condensed chromosomes attach to the spindle via a specialized region of chromatin called the centromere, ensuring equal partitioning of DNA into daughter cells. Centromeres are defined by the centromeric histone variant, CenH3, which is the foundational centromeric protein in most eukaryotes (Sullivan, Hechenberger et al. 1994, Yoda, Ando et al. 2000). First identified as Cenp-A in mammals (Earnshaw and Rothfield 1985, Palmer, O’Day et al. 1991), CenH3 localizes to centromeric DNA and helps recruit other components of the kinetochore, which mediates chromosome segregation. The loss of CenH3 results in catastrophic chromosome segregation defects and lethality in protists, yeast, flies, nematodes, mice, and plants (Stoler, Keith et al. 1995, Buchwitz, Ahmad et al. 1999, Howman, Fowler et al. 2000, Blower and Karpen 2001). Although some lineages lack CenH3 altogether (Akiyoshi and Gull 2014, Drinnenberg, deYoung et al. 2014), in most eukaryotes that encode CenH3, it is essential for chromosome segregation in both mitosis and meiosis.

In addition to CenH3’s critical role in mitotic and meiotic chromosome segregation, CenH3 protein retention is important for the epigenetic inheritance of centromere identity through spermiogenesis. During the production of male gametes in many animal species, the sperm nucleus undergoes a dramatic transition from histone-based chromatin to chromatin that is packaged by protamines; nearly all of the histones are removed and are replaced by highly basic proteins called protamines (Oliva and Dixon 1991, Braun 2001, Renkawitz-Pohl, Hempel et al. 2005). Even though CenH3 is a histone protein, it is not removed from sperm chromatin during this process. Studies in mammals find the presence of CenH3 in mature sperm (Palmer, O’Day et al. 1990). Furthermore, loss of paternal CenH3 on sperm chromatin in *Drosophila melanogaster* results in early embryonic lethality (Raychaudhuri, Dubruille et al. 2012). Thus, CenH3 needs to function in disparate chromatin environments in multicellular animals, in a histone-rich environment in somatic cells and in a protamine-rich environment in sperm, which may impose divergent functional constraints on CenH3.

The female germline could also impose distinct constraints on CenH3 function, particularly in long-lived animals. In humans and mice, oocyte nuclei arrest in meiotic prophase I for extended periods of time (years in humans, months in mice) (Von Stetina and Orr-Weaver 2011, Smoak, Stein et al. 2016). Oocyte centromere function does not seem to depend on the loading of newly transcribed CenH3 as conditional knockouts of CenH3 in meiotic prophase I are fully fertile in *Mus musculus* (Smoak, Stein et al. 2016). However, recent work demonstrated that CenH3 in MI arrested starfish oocytes undergoes gradual turnover, presumably to replace CenH3 containing nucleosomes that are disturbed by transcriptional machinery, allowing oocytes to maintain centromere competence over long periods of time (Swartz, Mckay et al. 2018). This means that CenH3 molecules are capable of stably persisting in oocytes for long periods of time and that there are mechanisms in place to maintain centromere function in non-dividing cells.

These separate functional requirements could impose opposite selective constraints on CenH3. For instance, one might anticipate that CenH3’s essential mitotic function would lead to functional constraint and strong amino acid conservation (‘purifying selection’). Contrary to this expectation, CenH3 has been found to evolve rapidly in many species of plants and animals (Malik and Henikoff 2001, Talbert, Bryson et al. 2004, Schueler, Swanson et al. 2010). We previously hypothesized that this rapid evolution is a result of CenH3’s role as a suppressor of centromere drive, which results from an inherently asymmetric transmission of chromosomes through female meiosis in both plants and animals (Malik 2009, Kursel and Malik 2018).

Because of these disparate functions, CenH3 proteins may have different protein coding requirements in different cellular contexts, especially in the germline (Das, Smoak et al. 2017). Dissecting these multiple functional constraints in many model organisms (such as *D. melanogaster* and *M. musculus*) are challenging because CenH3 is an essential single copy gene in these species. However, organisms in which CenH3 has duplicated and may have partitioned these specialized functions among paralogs present a unique opportunity to more precisely understand the requirements of CenH3 in each specific cellular context. Indeed, some plant species have multiple CenH3 paralogs (Finseth, Dong et al. 2015, Maheshwari, Tan et al. 2015) that show signs of tissue-specific specialization. For example, knockdown of one CenH3 paralog in wheat causes growth defects whereas knockdown of the other paralog causes reproductive defects (Yuan, Guo et al. 2015). However, the molecular basis of this specialization is unclear.

In contrast to plants, centromeric histone specialization has not been previously observed in animal species. Although an estimated 10% of plant genomes harbor multiple CenH3 paralogs (Kawabe, Nasuda et al. 2006, Finseth, Dong et al. 2015, Maheshwari, Tan et al. 2015), CenH3 duplications were previously thought to be rare in animals (Li and Huang 2008, Monen, Hattersley et al. 2015). Contrary to this view, we recently found that the CenH3 gene in Drosophila (known as *Cid*) has duplicated at least four times (Kursel and Malik 2017). Our evolutionary analyses revealed that thousands of Drosophila species (likely the majority of known Drosophila species) encode more than one *Cid* gene. We found that all *Cid* paralogs can localize to centromeres when ectopically expressed, but many paralogs have evolved germline-restricted expression patterns, highly divergent N-terminal tails and divergent selective constraints. This discovery led us to hypothesize that Drosophila Cid paralogs have acquired tissue or cell-type-specific functions (Kursel and Malik 2017).

To test this hypothesis, we performed cytological analysis of the two *Cid* paralogs in *Drosophila virilis, Cid1* and *Cid5*, which diverged nearly 40 million years ago and have since been co-retained in the Drosophila subgenus (Kursel and Malik 2017). We examined Cid1 and Cid5 localization in *D. virilis* somatic cells, testes, ovaries and early embryos. We found that there is mutually exclusive retention of the two Cid proteins in mature male and female gametes, which is achieved by alternate protein loss during meiosis in males and females. We hypothesize that paralog-specific changes in the N-terminal domain have allowed for the functional specialization of Cid1 and Cid5. Thus, Cid paralogs in *D. virilis* appear to have used gene duplication and specialization to resolve the tension of multiple, disparate CenH3 functions. This specialization further suggests that single copy CenH3 proteins may not represent the most optimal state in multicellular, sexual organisms.

## Results

The ancient retention of *Cid1* and *Cid5* suggests that both paralogs perform important, non-redundant, functions (Kursel and Malik 2017). In order to gain insight into the function of Cid1 and Cid5, we investigated their localization in dividing somatic cells, ovaries, testes and embryos of *D. virilis* flies. For this approach, we developed tools to visualize Cid1 and Cid5 *in vivo*. We exploited the high divergence of their N-terminal tails to develop polyclonal antibodies that are specific to either Cid1 or Cid5 (Figure S1A). We confirmed that each antibody specifically recognized the paralog it was designed for, in immunofluorescence analyses (Figures S1B and S1C).

Since antibody occlusion could hamper cytological analyses especially in the male germline (Bonnefoy, Orsi et al. 2007), we also generated transgenic *D. virilis* flies with *Cid1GFP* or *Cid5mCherry* under the control of their respective native promoters. In *D. melanogaster, Cid-GFP* transgenic flies, in which GFP was inserted between the N-terminal tail and histone fold domain of *Cid*, can complement *Cid* function (Schuh, Lehner et al. 2007). Therefore, we inserted the fluorescent protein tag between the N-terminal tail and the histone fold domain in both Cid1 and Cid5 transgenes (Figure S1D).

### Cid1, but not Cid5, is detectable in somatic cells

Our previous expression analyses based on RT-PCR (Kursel and Malik 2017) found that *Cid1* is expressed in somatic cells including *D. virilis* WR-Dv-1 tissue culture cells (derived from first instar larvae), heads from male and female *D. virilis* flies and male and female carcasses (decapitated, gonad-ectomized, animals), whereas *Cid5* is not. To examine protein expression, we looked for Cid1 and Cid5 protein in two types of dividing somatic cells: tissue culture cells and larval neuroblasts. In WR-Dv-1 cells, we could detect endogenous Cid1 protein by both western blot and immunofluorescence analyses (Figure 1A, Figure 1B). However, we did not detect Cid5 using either method (Figure 1A, Figure 1C), consistent with our previous finding that *Cid5* RNA is not found in these cells (Kursel and Malik 2017).

**Figure 1.**
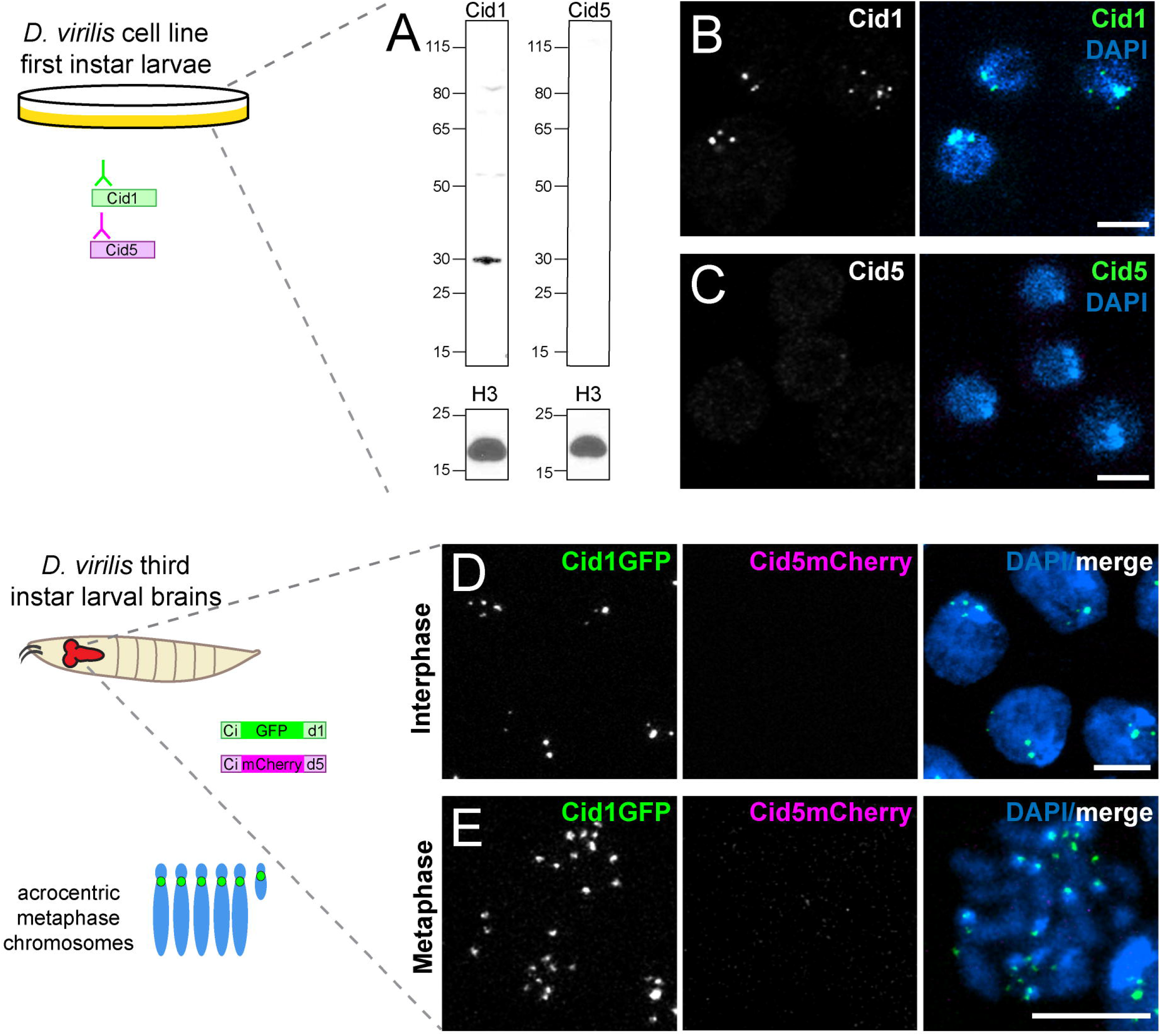
Cid1 is the centromeric histone in two dividing somatic cell types. (A) Western blot of Cid1 and Cid5 in *D. virilis* tissue culture cells. A western blot for histone H3 was used as a loading control. This western blot was repeated three times with the same result. (B - C) Immunofluorescence images of *D. virilis* tissue culture cells stained with Cid1 (B) or Cid5 (C) antibodies. (D - E) Images of interphase (D) or metaphase (E) cells from *D. virilis* larval brains dissected from flies containing Cid1GFP and Cid5mCherry transgenes. Scale bar = 5μm.

Next, we examined Cid1 and Cid5 localization in larval neuroblasts, a tissue that is enriched in mitotic cells. As expected, we found that Cid1 localized to centromeres in interphase cells and on condensed metaphase chromosomes (Figure 1D, Figure 1E). As *D. virilis* chromosomes are acrocentric (have their centromeres close to one telomere) the Cid1 signal was localized to one end of each condensed chromosome. In contrast, we could not detect any Cid5 signal (Figure 1D, Figure 1E). Our cytological findings using transgenes were confirmed by detection using polyclonal antibodies, reinforcing the validity of our transgene analyses (Figure S1B).

### Differential localization of Cid1 and Cid5 in D. virilis ovaries

We next investigated Cid1 and Cid5 protein localization in *D. virilis* ovaries. The *Drosophila* ovary is made up of about 16 ovarioles. At the anterior tip of each ovariole, germline stem cells divide four times to produce a cyst of 16 interconnected cells, which differentiate into 15 nurse cells (support cells that provide mRNA, protein and other material to the oocyte via a shared cytoplasm) and one oocyte. These interconnected germ cells are surrounded by somatic follicle cells and together form an egg chamber. Egg chamber maturation occurs progressively along the ovariole in a series of defined stages. These stages are referred to as stages 1-14 based on growth and organization of somatic and germline cells. By stage 2, the oocyte has entered into meiotic prophase and reaches pachytene. At stage 5, the oocyte enters primary arrest and remains arrested until stage 13 when the oocyte progresses to secondary arrest in metaphase of meiosis I. In the stage 14 egg chamber, no nurse cell nuclei remain and the oocyte is prepared for ovulation (King, Rubinson et al. 1956, Spradling 1993). Our previous study showed that *Cid1* transcripts were abundant but *Cid5* transcripts were not detectable in RNA extracted from whole ovaries (Kursel and Malik 2017). We, therefore, expected to find that Cid1 would be the only Cid paralog detectable in *D. virilis* ovaries.

To examine Cid1 and Cid5 protein, we used Cid1GFP and Cid5mCherry transgenic flies and Cid1 and Cid5 antibodies for localization of both proteins in somatic and germline cells at different stages of egg chamber development. Similar to mitotically dividing somatic cells, we detected Cid1 but not Cid5 in somatic follicular cells (Figure 2A, Figure 2B). However, we were surprised to find that both Cid1 and Cid5 protein were robustly detected in the germline lineage cells of the ovary in egg chamber stages 2 – 9 (Figure 2A), including in nurse cells (Figure 2C) and the oocyte nucleus (Figure 2D). We similarly detected Cid1 and Cid5 protein in germline cells at these stages using the Cid1 and Cid5 antibodies (Figure S2). However, by stage 14, Cid1 was the only detectable paralog at centromeres of metaphase I arrested chromosomes (Figure 2E) via Cid-GFP visualization (compare Cid-GFP staining in Fig. S2G to Cid5-mCherry staining in Fig. S2H); however, antibody staining against Cid1 is not successful at this late stage of oogenesis (Fig. 2H) likely due to antibody accessibility, highlighting the utility of our dual approaches.

**Figure 2.**
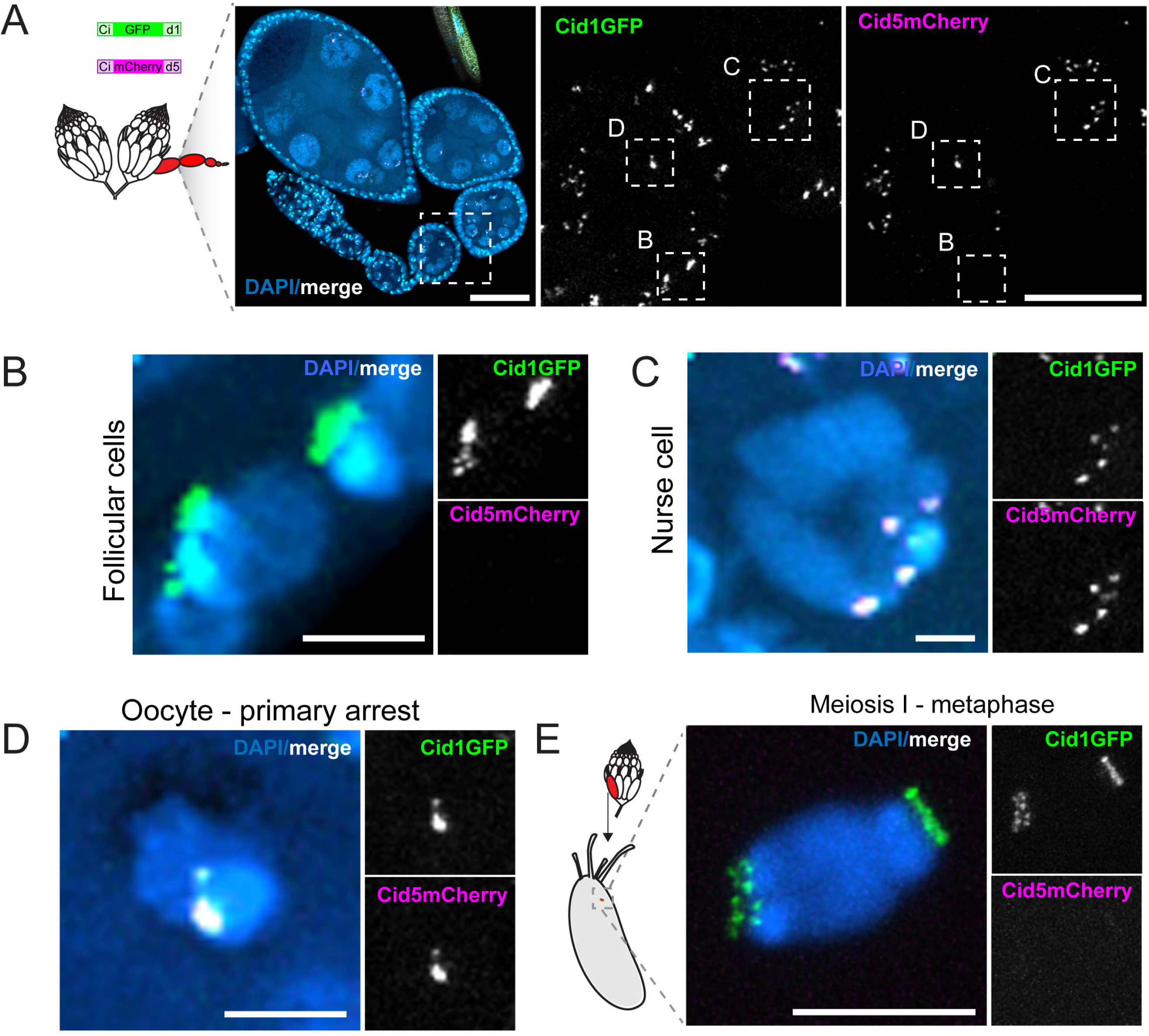
Differential localization of Cid1 and Cid5 in ovaries. All images use Cid1GFP and Cid5mCherry to detect Cid1 and Cid5 protein. (A) A whole *D. virilis* ovariole. The two right panels show the boxed region of the left panel at higher magnification. Regions surrounded by boxes in the right two panels are shown at high magnification in subsequent panels. Scale bar = 45μm in left panel, 20μm in the right panel. (B) High magnification image of follicular cells (somatic) from a stage 3 egg chamber. (C) High magnification image of a nurse cell from a stage 4 egg chamber. (D) High magnification image of a stage 3 oocyte in primary arrest. (E) Image of a stage 14 oocyte nucleus in meiosis I metaphase arrest. Scale bars in (B) – (E) = 5μm.

These results suggest that both Cid1 and Cid5 are present at centromeres early in oogenesis but only Cid1 remains on centromeres by meiosis I metaphase arrest. Given that turnover of CenH3-containing nucleosomes in MI-arrested oocytes appears to be quite gradual (~2% of centromeric CenH3 is exchanged per day in MI-arrested starfish oocytes (Swartz, Mckay et al. 2018)), we hypothesize that Cid5 protein is actively removed from the oocyte centromeres and at the onset of meiosis I metaphase arrest. Since Cid1 is always present throughout oogenesis, it is unclear whether Cid5 performs any function, centromeric or otherwise, in the female germline. However, it is apparent that Cid1 is the only detectable centromeric histone in late-stage *D. virilis* oocytes and is therefore likely to be essential for female fertility and early embryonic mitotic divisions following fertilization.

### Differential localization of Cid1 and Cid5 in D. virilis testes

Our previous characterization of *Cid1* and *Cid5* mRNA expression in *D. virilis* (Kursel and Malik 2017) indicated that both *Cid* paralogs are expressed in testes. We therefore examined the cytological localization patterns of Cid1 and Cid5 in the *D. virilis* male germline. In the *Drosophila* male germline, spermatogenesis begins at the apical tip of the testis where the germline stem cells reside. The asymmetric divisions of the germline stem cells replenish the stem cell population and produce gonialblasts. These gonialblasts divide mitotically with incomplete cytokinesis and then enter an extended meiotic prophase. Following this extended period of cell growth, cysts of 16 spermatocytes undergo meiosis and produce bundles of 64 haploid spermatids (Fuller 1993, Fabian and Brill 2012). These spermatids then go through the process of nuclear remodeling resulting in 200-fold compaction of their nuclear volume (Fuller 1993). During this dramatic nuclear reorganization, nearly all of the histones are removed and are replaced by sperm nuclear basic proteins (SNBPs) (Renkawitz-Pohl, Hempel et al. 2005). Finally, elongated spermatid bundles go through individualization to produce mature sperm (Figure 3A, Figure 3B).

**Figure 3.**
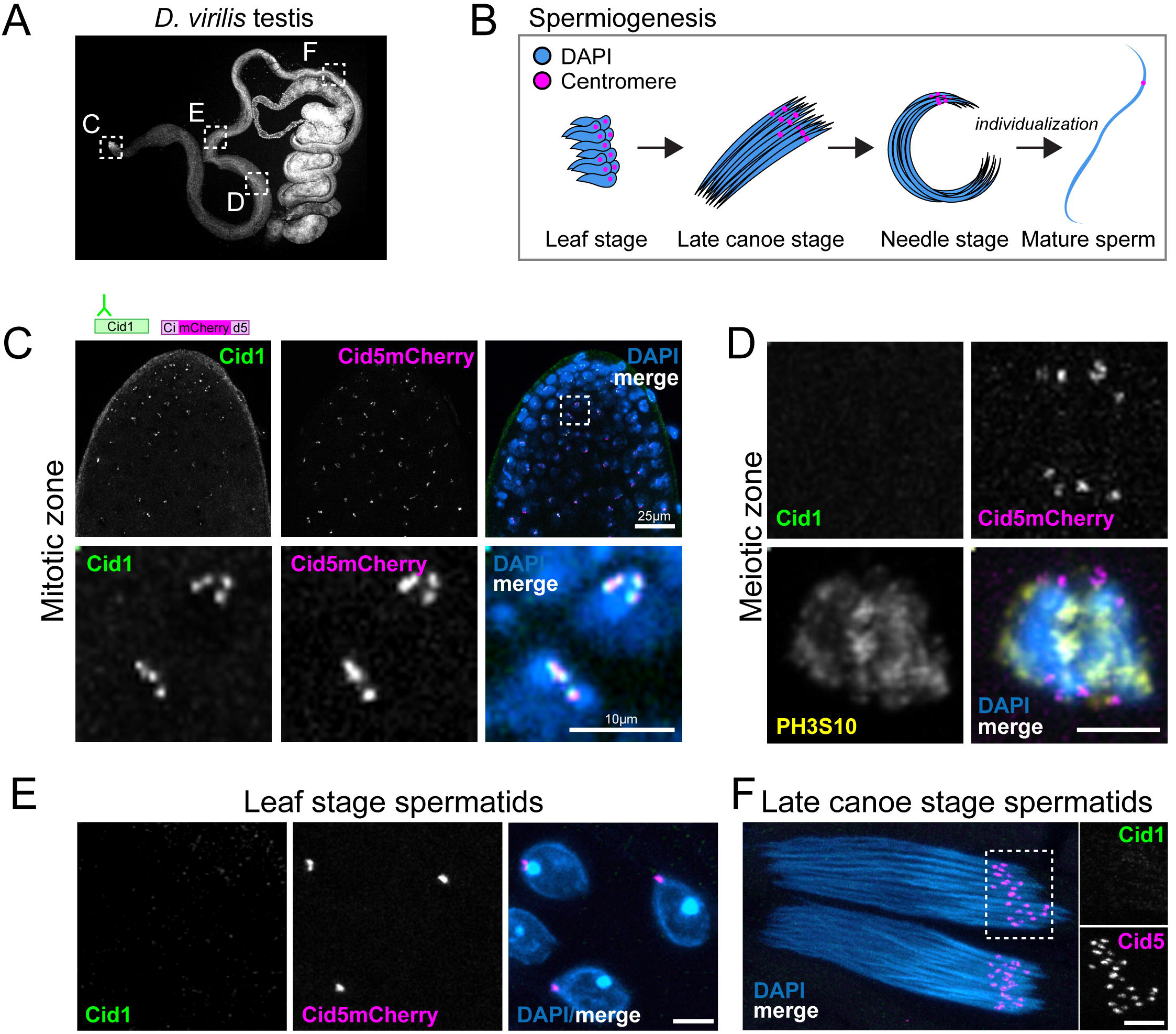
Differential localization of Cid1 and Cid5 in testes. (A) Image of a DAPI stained *D. virilis* testis. Boxed regions show the approximate location of panels (C) – (F). Scale bar = 100μm. (B) Schematic showing stages of spermiogenesis. The Cid1 antibody and Cid5mCherry transgene were used to visualize Cid1 and Cid5 in the images in (C) – (F). (C) The apical tip (mitotic zone) of a *D. virilis* testis. The bottom panel shows a high magnification image of the area indicated in the top panel by the dashed box. Scale bar = 25μm in top panel and 10μm in the bottom panel. (D) A single cell with condensing chromosomes in late prometaphase or early metaphase. PH3S10 antibody staining is also shown. (E) Leaf-stage spermatid nuclei. (F) Late-canoe stage spermatid bundles. Scale bars = 5 μm in (D) – (F).

Previous studies in *D. melanogaster* have shown that Cid is essential for the mitotic and meiotic divisions in the male germline (Dunleavy, Beier et al. 2012). Moreover, Cid has also been shown to be critical for transgenerational centromere inheritance through the mature sperm (Raychaudhuri, Dubruille et al. 2012). Therefore, we examined the cytological localization of Cid1 and Cid5 in the mitotic zone, meiotic zone, post-meiotic stages and in mature sperm (Figure 3A, Figure 3B). We examined testes from Cid5mCherry males and performed antibody staining with the Cid1 antibody and a phospho-histone H3 Serine 10 (PH3S10) antibody to identify condensed chromosomes (Hendzel, Wei et al. 1997, Tang, Bickel et al. 1998, Ivanovska and Orr-Weaver 2006). We found that Cid1 and Cid5 co-localize at centromeres in the mitotic zone of the testis (Figure 3C). However, surprisingly, at the onset of metaphase of meiosis I, Cid1 was no longer observed, and we could only detect Cid5 on these chromosomes (Figure 3D). We could also detect Cid5 in post-meiotic stages as a discrete focus on each ‘leaf-stage’ and ‘late-canoe-stage’ spermatid nucleus (Fabian and Brill 2012), but we never observed Cid1 at these stages (Figure 3E, Figure 3F).

To confirm that our inability to detect Cid1 in meiotic cells and post-meiotic spermatids was not due to antibody accessibility issues, we also examined Cid1GFP in the male germline. The results were nearly identical to the antibody staining. We detected Cid1 at centromeric foci in the mitotic zone (Figure S3A) but we could only detect faint Cid1 signal in cells entering metaphase of meiosis I zone (Figure S3B). We could not detect Cid1 at any stage after meiosis, including in mature sperm (Figure S3C – S3E). Our results are thus consistent between our antibody staining and transgene analyses, except for cells entering meiosis I metaphase, in which Cid1GFP is either slightly more sensitive than the Cid1 antibody, or persists longer than endogenous Cid1. Regardless, these results indicate that metaphase of meiosis I represents a transition state between the presence of Cid1 in mitotic and early meiotic cells and its absence in post-meiotic cells. Like the loss of Cid5 in oocytes, this loss of Cid1 occurs without DNA replication, suggesting an active protein degradation mechanism may be responsible. Interestingly, previous studies in *D. melanogaster* testes also observed a decrease in Cid levels coinciding with changes in kinetochore organization and orientation between meiosis I and meiosis II (Dunleavy, Beier et al. 2012). Thus, metaphase of meiosis I represents a centromeric transition state in both males and females, except that Cid5 is specifically lost in the female germline and Cid1 is specifically lost in the male germline.

Our cytological analyses further indicate that Cid5’s centromeric localization persists throughout male gametogenesis from early germ cells to sperm. Previous findings have demonstrated that Cid protein is required for transgenerational inheritance of centromere identity through sperm in *D. melanogaster* (Raychaudhuri, Dubruille et al. 2012). Since Cid1 is not detectable during spermiogenesis, we hypothesized that Cid5 might provide the transgenerational centromeric mark in mature sperm in *D. virilis*. To further investigate Cid5 localization in *D. virilis* sperm and validate its centromeric localization, we employed GFP-Hiphop as a telomeric (and therefore centromere-adjacent) marker (Gao, Cheng et al. 2011). Since *D. virilis* flies have acrocentric chromosomes, their centromeric cytological signals should be adjacent to one of the two telomeric, GFP-Hiphop-labeled, cytological signals on each chromosome. Thus, Hiphop localization serves as an additional centromere-adjacent marker in *D. virilis*.

We examined the localization of GFP-HipHop and Cid5mCherry in the testes of flies that contained both transgenes. We observed two primary HipHop foci corresponding to telomeric ends in each spermatid nucleus (Figure 4A, 4B). We also saw a single Cid5 focus, which consistently co-localized with one of the two GFP-HipHop foci (Figure 4C). This localization pattern persisted throughout spermatid development and in mature sperm (Figure 4C - 4E). These experiments give additional support to the hypothesis that Cid5 provides the transgenerational centromere mark in *D. virilis* – Cid5 is present at centromeres in mature sperm, but Cid1 is not.

**Figure 4.**
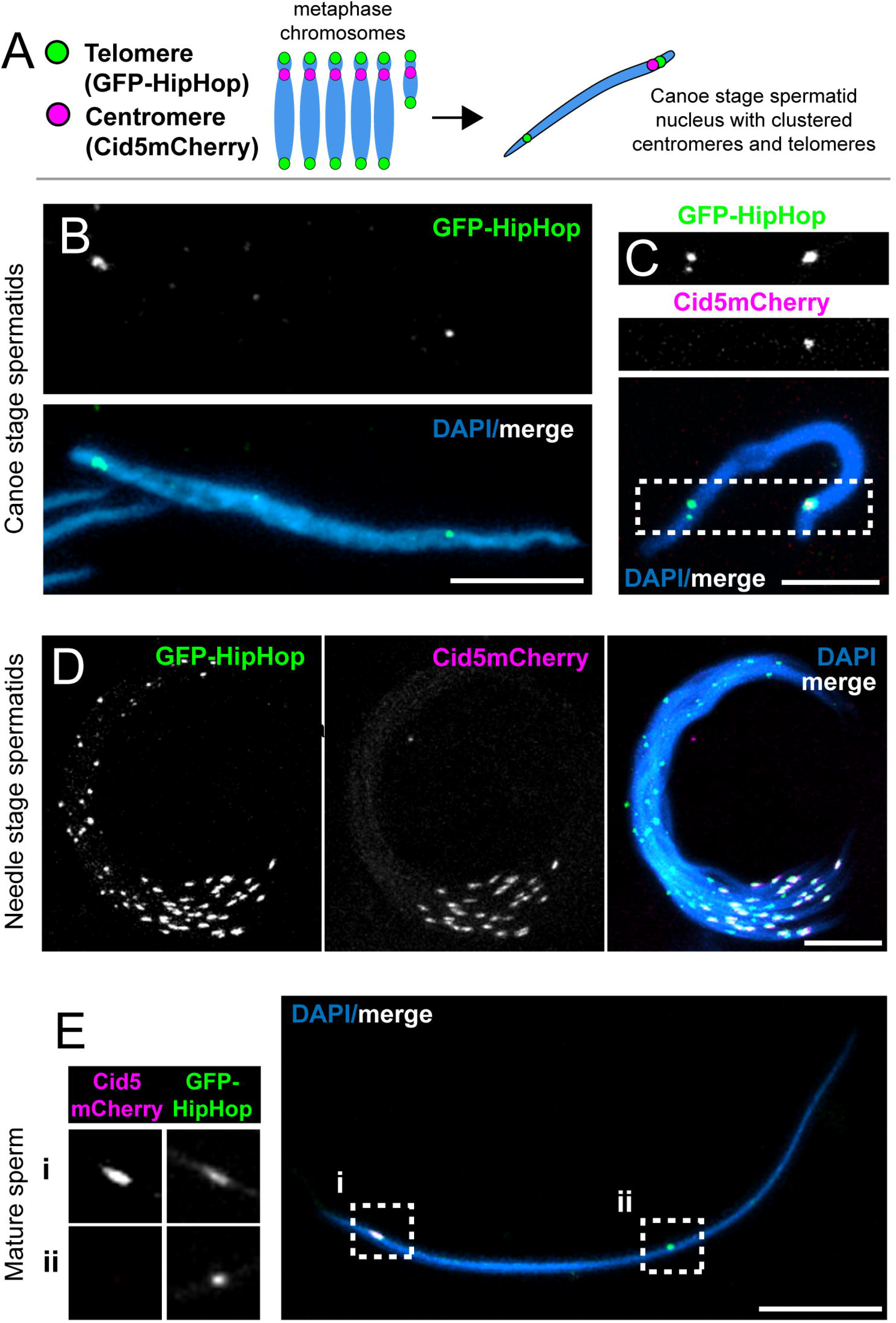
Cid5 provides the centromere mark in mature sperm. (A) Schematic showing haploid chromosomes (left) which become condensed into the Rabl configuration (right) in *D. virilis* sperm. (B) A single late canoe stage spermtid from a GFP-HipHop fly. All subsequent panels show images from flies with both GFP-HipHop and Cid5mCherry transgenes. (C) A single late-canoe stage spermatid. (D) Needle-stage spermatid bundle. (E) A single mature sperm nucleus. Boxed regions (i) and (ii) are also shown at slightly higher magnification and as separate channels (left). All scale bars = 10μm.

Taken together, our cytological examination of Cid1 and Cid5 in the *D. virilis* male germline suggests that after prometaphase, Cid5 is the predominant centromeric histone. Our inability to robustly detect Cid1 in post-prometaphase meiotic cells and post-meiotic spermatids strongly suggests that male meiotic and centromere inheritance function in *D. virilis* flies does not require Cid1, even though the *D. melanogaster Cid1* ortholog, *Cid*, is essential for both processes (Dunleavy, Beier et al. 2012, Raychaudhuri, Dubruille et al. 2012). Thus, male and female gametes alternately retain different Cid protein paralogs in *D. virilis*.

### Maternal Cid1 rapidly replaces paternal Cid5 following fertilization

Our cytological analyses revealed that the mature oocyte nucleus in *D. virilis* only retains Cid1 whereas mature sperm only retain Cid5 (Figure 5A). We next investigated how parental genomes with distinct Cid paralogs coordinate chromosomal events in the early embryo. One of the most dramatic chromosomal changes following fertilization is the remodeling of the sperm nucleus, in which SNBPs (protamines in mammals), which package the bulk of sperm chromatin, are replaced with maternally-provided core and variant histones in a replication-independent manner (Loppin, Docquier et al. 2000, Loppin, Berger et al. 2001, Loppin, Bonnefoy et al. 2005). In *D. melanogaster*, paternal Cid persists on the paternal genome throughout this extensive remodeling and is required for the first embryonic cell divisions, even though the specific molecules of paternal Cid only persist until the third embryonic cell cycle (Raychaudhuri, Dubruille et al. 2012). While paternal chromosome remodeling occurs, female meiosis is completed. Maternal and paternal pronuclei then congress towards each other, appose and undergo mitosis synchronously but on separate halves of the first spindle (Figure 5A). Defects in this synchronization lead to embryonic lethality (Landmann, Orsi et al. 2009, Levine, Vander Wende et al. 2015).

**Figure 5.**
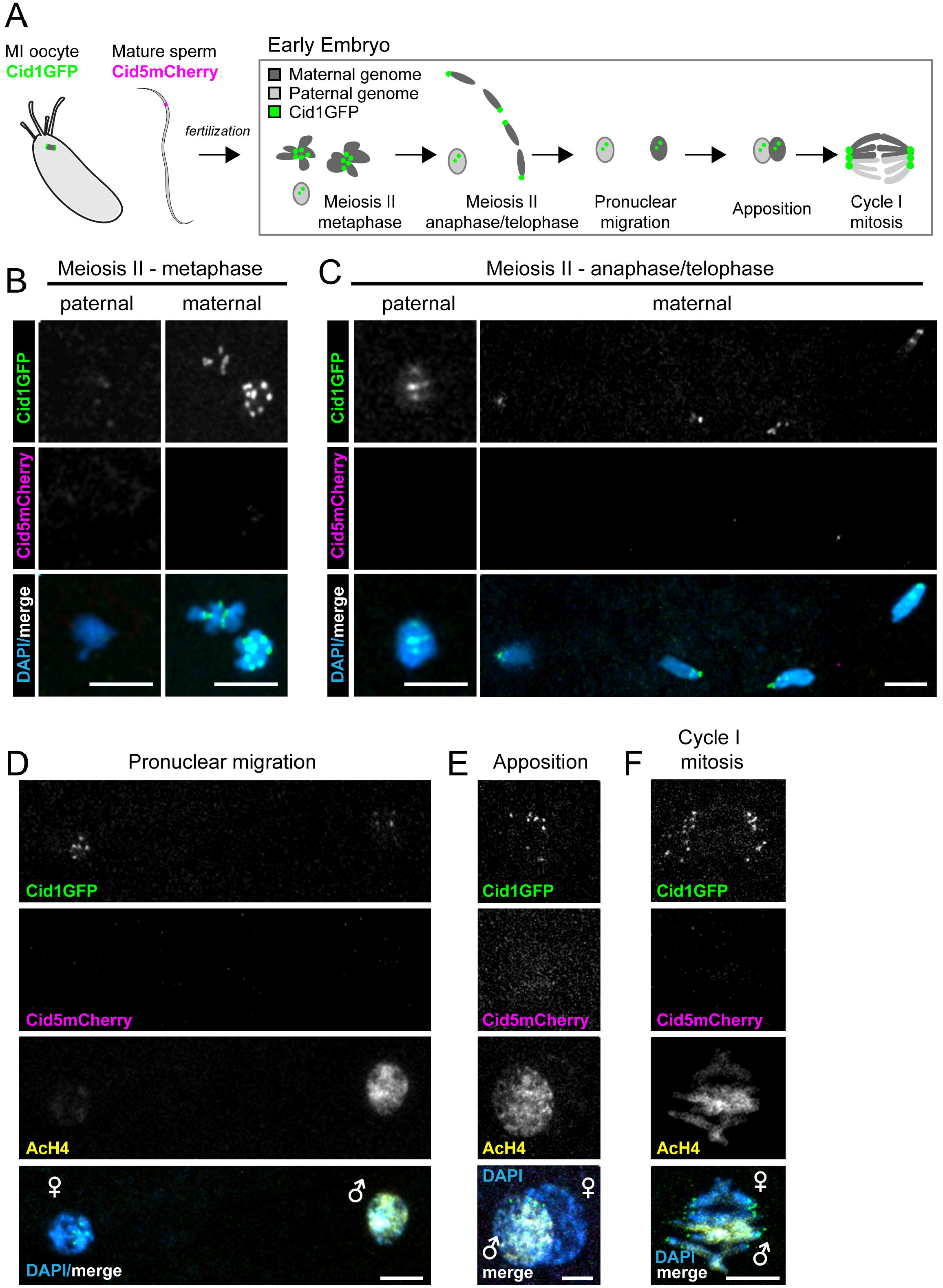
Cid1 replaces Cid5 in the early embryo. (A) Schematic of fertilization and the progression of the maternal and paternal genome in the early embryo through the first embryonic mitosis. All other panels are images from *D. virilis* early embryos that were collected from parents that both had Cid1GFP and Cid5mCherry transgenes. Paternal and maternal genomes were discerned by nuclear morphology, (B) and (C), or by acetylated histone H4 (AcH4) antibody staining, (D) – (F). AcH4 preferentially stains the paternal genome. (B) Meiosis II metaphase. (C) Meiosis II anaphase/telophase. (D) – (F) Pronuclear migration, apposition and the first embryonic mitotic cell division. All scale bars = 5μm.

Based on the precedent in *D. melanogaster*, we expected that paternally inherited Cid5 would persist on the paternal genome through the first several embryonic cell cycles, whereas Cid1 would define centromeres throughout the completion of female meiosis, co-localize with Cid5 in the early embryo and gradually replace Cid5 to eventually become the only Cid protein present in the embryo. To test this hypothesis, we examined Cid1GFP and Cid5mCherry in embryos produced by male and female parents bearing both transgenes. Consistent with our previous findings that meiosis I metaphase arrested oocytes only contain Cid1 (Figure 2E), we found that only Cid1 is detectable on the maternal genome through the completion of meiosis (Figure 5B, Figure 5C). More surprisingly, we were only able to only detect Cid1 on the paternal pronucleus, even at very early stages (Figure 5B, Figure 5C) despite our earlier observations that mature sperm only contain readily detectable Cid5 (Figure 3E, 4E). Although Cid1 signal was faint on the paternal genome at earlier stages, it rapidly reached a level comparable to the Cid1 signal on the maternal genome by the time of the synchronous first mitosis (Figure 5B - 5F). Our results suggest that in *D. virilis*, maternal Cid1 replaces paternal Cid5 even more rapidly than it does in *D. melanogaster*, during the protamine-to-histone chromatin transition prior to the first mitotic division.

## Discussion

Our study reveals that *D. virilis* employs a dedicated CenH3 paralog, Cid5, specifically for the purpose of epigenetic inheritance of centromere identity through sperm. This specialization was accomplished subsequent to the gene duplication event that birthed Cid5 by virtue of gain of germline-specific expression of Cid5, specific removal of the ancestral Cid1 during male meiosis prior to spermiogenesis, and replacement of Cid5 with Cid1 following fertilization. Intriguingly, a reduction in Cid protein levels is also observed in *D. melanogaster* between the two meiotic divisions in the male germline (Dunleavy, Beier et al. 2012). However, unlike in *D. virilis*, in *D. melanogaster* it is essential that Cid (the Cid1 homolog) persists through meiosis and is present in mature sperm (Raychaudhuri, Dubruille et al. 2012). Although Cid1 no longer appears to perform centromere inheritance function through the male germline in *D. virilis*, it still serves CenH3 function in both the soma and the female germline.

Based on this, we predict that *D. virilis* flies lacking Cid1 would be inviable, just like *Cid* knockdown *in D. melanogaster* (Blower and Karpen 2001) whereas Cid5 knockouts would result in either male sterility or paternal effect lethality as observed in *D. melanogaster* when Cid is specifically depleted in sperm (Dunleavy, Beier et al. 2012, Raychaudhuri, Dubruille et al. 2012). However, it is also possible that Cid1 is required for loading Cid5 onto centromeres in male germ cells, in which case Cid1 knockdown in the male germline would also result in sterility. Nevertheless, we anticipate that due to their specialization, knockout or genetic knockdown of *Cid1* and *Cid5* will have different phenotypic consequences, just as previously observed for wheat *CenH3* paralogs (Yuan, Guo et al. 2015).

What is the molecular basis of the specialization of Cid paralogs in *D. virilis*? Although Cid1 and Cid5 have diverged in their histone fold domains (HFDs), we speculate that the primary mode of specialization is via the much greater divergence of their N-terminal tail domains (NTDs). Cid1 and Cid5 differ significantly in both the retention of ancestral conserved motifs and in their acquisition of new motifs in their NTDs. All single copy *Cid* genes in *Drosophila* (including from *D. melanogaster*) encode a highly stereotyped set of protein sequence motifs 1 – 4 in their NTDs (Malik, Vermaak et al. 2002, Kursel and Malik 2017), which are also conserved in *D. virilis* Cid1. Motifs 1 – 3 have been implicated in sister centromere cohesion in male meiosis (Collins, Malacrida et al. 2018) whereas motif 4 has been associated with BubR1 recruitment (Torras-Llort, Medina-Giro et al. 2010). However, Cid5 proteins have lost motifs 1 and 3 (Kursel and Malik 2017), suggesting that these motifs are not necessary for Cid5’s role in male meiosis and spermiogenesis. Differences in Cid1 and Cid5’s NTDs could result in different protein interactions and cell-type specific kinetochore formation. Furthermore, both Cid1 and Cid5 have gained new motifs (motif 8 in Cid1, motifs 9 and 10 in Cid 5) not found in any Cid proteins encoded by single copy genes (*e.g*., Cid in *D. melanogaster*) (Kursel and Malik 2017). We speculate that these ‘new’ motifs may represent degron domains that underlie the specific loss of Cid1 and Cid5 in male and female meiosis respectively in *D. virilis*. Thus, either motif would be highly deleterious if a single Cid protein encoded both male and female germline function.

*Cid* paralogs like the two we have described in *D. virilis* have allowed the gametic functions of CenH3 to be separated into different genes; this separation of function has since been refined over millions of years of natural selection. These paralogs allow us to develop concrete hypotheses regarding the specific functional roles of N-terminal motifs that would not be otherwise possible in species carrying only one essential copy of *CenH3*. At least two other ancient *Cid* duplication events have been found in the montium group of *Drosophila* species. Some of these paralogs also show testis-specific expression patterns, just like in *D. virilis* (Kursel and Malik 2017). Moreover, their N-terminal tails show a similar pattern of gain and loss of motifs. Investigations in these species will reveal whether the evolution of *Cid* paralogs follows a convergent trajectory of functional specialization.

Our study of Cid paralogs in *D. virilis* reveals that Cid1 and Cid5 carry out distinct roles in different cell types. Given the disparate nature of the chromatin environment in which each paralog functions, we propose that Cid1 and Cid5 face different selective pressures. Moreover, we propose that single copy *CenH3* genes must encode all of the functions performed by Cid1 and Cid5. If these roles are equally important for fitness, a single *CenH3* gene encoding both functions could become ‘trapped’ for suboptimal function in both roles *e.g*., soma versus sperm. Such ‘intralocus conflict’ occurs in the case of sexual genetic conflicts, whereby a locus beneficial in one sex is detrimental to the other (VanKuren and Long 2018). However, the same functional tradeoff might also result if a single gene had two functional optima that could not be simultaneously achievable. One way to resolve intralocus conflict is through gene duplication and specialization of different paralogs for different functions (Gallach and Betran 2011). For example, genes encoding mitochondrial function may have divergent optima in the soma versus testis, and this divergence has been invoked to explain the high retention rate of testis-specific gene paralogs encoding mitochondrial function (Gallach, Chandrasekaran et al. 2010). If CenH3 function is also subject to dual constraints, then the duplication and specialization of different Cid paralogs in species like *D. virilis* may represent a more optimal state than the single copy *Cid* gene in species like *D. melanogaster*. Under this scenario, the Cid1 and Cid5 paralogs of *D. virilis* provide an elegant example of nature’s ‘separation-of-function’ experiment for a CenH3 gene that has multiple essential functions in multicellular organisms.

## Materials and methods

### Cid1 and Cid5 antibody production

We raised an antibody against Cid1 residues 15 – 31 (KSESHLDNVEDSYEKTA) and Cid5 residues 56 - 71 (NLESPVAGEEPAPDTV). These sites were selected because they are in regions where Cid1 and Cid5 share no apparent homology and are distinct from other *D. virilis* proteins. Covance Inc. (Princeton, NJ) immunized two rabbits with the conjugated Cid5 peptide by injecting it four times over the course of four months. Covance also immunized two rabbits for the Cid1 peptide by injecting it five times over the course of five months. Our previous analysis of *D. virilis* Cid5 polymorphism revealed non-synonymous variation in the Cid5 peptide sequence used to generate the antibody, therefore, for all experiments we ensured that we used *D. virilis* strains and cell lines have appropriate Cid5 alleles.

### Western blots from *D. virilis* WR DV-1 cells

*D. virilis* WR DV-1 cells were collected in RIPA buffer and sonicated. Protein was quantified by Bradford assay and 20ug total protein was analyzed by western blot. We probed the membrane with either rabbit anti-Cid1 (1:2000), rabbit anti-Cid5 (1:2000), or rabbit anti-H3 (1:5000 Abcam ab1791) primary antibodies followed by goat anti-rabbit IgG-HRP (1:5000 Santa Cruz Biotechnologies Inc., Dallas, TX).

### Antibody staining of *D. virilis* tissue culture cells

We confirmed that the Cid1 antibody works for cytology by immunostaining *D. virilis* WR DV-1 cells. Cells were transferred to coverslips and fixed in 4% PFA for 5 min and blocked with PBSTx (0.3% Triton) plus 3% BSA for 30 minutes at room temperature. Then cells were incubated with primary antibodies at 4°C overnight. Coverslips with cells were incubated with secondary antibodies for 1 hour at room temperature. Antibodies were diluted as follows: rabbit anti-Cid1 1:5000 and (Invitrogen Alexa Fluor 568 A-11011) 1:2000.

### Overexpression of Venus-Cid5 in *D. melanogaster* KC cells

Since *Cid5* is not expressed in *D. virilis* tissue culture cells, we confirmed that the Cid5 antibody works for cytology by overexpressing Venus-Cid5 in *D. melanogaster* KC cells and performing immunostaining. Venus-Cid5 was cloned into an expression vector from the Drosophila Gateway Collection generating an N-terminal Venus (pHVW) fusion protein under the control of the *D. melanogaster* heat shock promoter. Transfections and antibody staining were performed as follows: two micrograms plasmid DNA was transfected using Xtremegene HP transfection reagent (Roche) according to the manufacturer’s instructions. 24 hours after transfection, cells were heat-shocked at 37°C for one hour to induce expression of the *Cid* fusion protein. Cells were transferred to a glass coverslip 24 hours after heatshock. Cells were fixed in 4% PFA for 5 min and blocked with PBSTx (0.3% Triton) plus 3% BSA for 30 minutes at room temperature. Coverslips were then incubated with primary antibodies at 4°C overnight. Coverslips with cells were incubated with secondary antibodies for 1 hour at room temperature. Antibodies were diluted as follows: rabbit anti-Cid51:2500 and goat anti-rabbit (Invitrogen Alexa Fluor 568, A-11011) 1:2000

### *D. virilis* transgenics

Cid1GFP and Cid5mCherry were cloned into a vector backbone containing piggyBac inverted repeats and the miniwhite gene cassette. This vector was generated by first removing 3XP3EGFP from the nosGal4-MW-pBacns plasmid (stock number 1290, Drosophila Genomics Resources Center). The 3XP3EGFP was removed as follows: nosGal4-MW-pBacns was digested with AgeI and AsiSI, run on a gel and the largest band was gel isolated. Overhangs were blunted, and then the vector was ligated to itself to produce nosGal4_MWonly. Then the nanosGal4 cassette was removed as follows: nosGal4_MWonly was digested with NotI, run on a gel and the largest band was gel isolated. Then the vector was ligated to itself to produce NoPromoter_miniwhite. Cid1GFP or Cid5mCherry, each with ~1kb sequence upstream and downstream, were inserted between the AvrII and SbfI sites of the NoPromoter_miniwhite plasmid. For both Cid1 and Cid5, fluorescent proteins were inserted immediately 5-prime of the RRRK motif at the beginning of the histone fold domain. Fluorophores were flanked on both sides by three glycine residues to function as flexible linkers. Cid1GFP and Cid5mCherry plasmids were injected along with the piggyBac helper plasmid phspBac (Handler and Harrell 1999) into *D. virilis* embryos. Injected flies were screened for red eye color. Injections and screening was performed by Rainbow Transgenics.

### Cytology: general data collection and presentation practices

For all cytological data, we present representative images acquired from the Leica TCS SP5 II confocal microscope with LASAF software and present maximally projected image files. For protein localization in larval neuroblasts, ovaries, testes and the early embryo, a minimum of five organs and five cell-types were examined for each assay of each stage.

### Preparation of larval neuroblasts for imaging and immunofluorescence

To assess Cid1 and Cid5 localization in larval brains, we used both Cid1 and Cid5 specific antibodies and Cid1GFP and Cid5mCherry transgenes. Brains from actively crawling third-instar larvae were dissected in PBS and transferred to 0.5% sodium citrate hypotonic solution 10 minutes. We transferred brains to a drop chromosome isolation buffer (120mg MgCl2:6H2O, 1g citric acid, 1mL Triton-X100, distilled H2O to 100mL) on a glass slide and fragmented the brains with needles for four minutes. Next, we lowered a coverslip was lowered onto the fragmented brains and squashed the brains under gentle pressure for 30 seconds. We then froze slides in liquid nitrogen. Then, slides were removed from liquid nitrogen and the cover slip was flipped off with a razor blade. Slides were immediately immersed in cold methanol for five minutes, cold acetone for one minute and PBS for one minute at room temperature. For experiments obtaining fluorescent signal from transgenes only, we removed the PBS and added mounting medium with DAPI. For antibody staining, after incubation in acetone, brains were rinsed once in PBS and then incubated in PBS + 1% TritonX for 10 minutes for permeabilization. Slides were blocked in PBS + 0.1% TritonX + 3% BSA for 30 minutes at room temperature. Slides were incubated with primary antibody overnight at 4°C. Then slides were washed and incubated with secondary antibodies for one hour at room temperature. Antibodies were diluted as follows: rabbit anti-Cid 1 1:1000, rabbit anti-Cid5 1:1000, and Alexa Fluor goat anti-rabbit 568 1:1000.

### Preparation of testes for imaging and immunofluorescence

To assess Cid1 and Cid5 localization in testes, we used Cid1 and Cid5 specific antibodies or transgenic flies encoding Cid1GFP or Cid5mCherry (both with internal tags and expressed under the control of their native promoters, as described above). To characterize Cid1 and Cid5 localization without antibody staining, we dissected testes in PBS from sexually mature (~10-day old) Cid1GFP, Cid5mCherry, or Cid1GFP/Cid5mCherry males. Testes were spread out on charged microscope slide, squashed under a coverslip and immediately immersed in liquid nitrogen. Testes were then fixed in 4% paraformaldehyde (PFA) for seven minutes or cold methanol (5 minutes) and acetone (5 minutes). Testes were then mounted in SlowFade Gold antifade with DAPI. For immunofluorescence, we fixed testes from Cid1GFP or Cid5mCherry transgenic flies in 4% PFA. Testes were permeabilized in PBS + 0.3% TritonX for 30 minutes (two-15 minutes washes) and blocked in PBS + 0.1% TritonX + 3% BSA for 30 minutes at room temperature. Primary antibodies were diluted in block and incubated with testes overnight. Secondary antibodies were incubated in block for one hour at room temperature. Antibodies were diluted as follows: mouse anti-phospho-histone H3 serine 10 (1:1000 Millipore clone 3H10) and Alexa Fluor goat anti-mouse 633 (1:1000).

### Preparation of ovaries for imaging and immunofluorescence

To assess Cid1 and Cid5 localization in ovaries, we used Cid1 and Cid5 specific antibodies or transgenic flies encoding Cid1GFP or Cid5mCherry (as described above). To characterize Cid1 and Cid5 localization without antibody staining, we dissected ovaries in PBS from sexually mature (~10-day old) Cid1GFP, Cid5mCherry, or Cid1GFP/Cid5mCherry *D. virilis* females. Ovaries were fixed in 1:1 paraPBT:heptane (paraPBT = 4% paraformaldehyde in PBS + 0.1% TritonX) for 10 minutes at room temperature. Then ovaries were washed, including one wash with 1X DAPI and mounted in SlowFade Gold (Thermo Fisher Scientific). For immunofluorescence, we performed fixation as above. We then blocked ovaries in PBS + 0.1% TritonX + 3% BSA for 30 minutes at room temperature. Ovaries were incubated with primary antibodies overnight at 4C. Ovaries were then washed and incubated with secondary antibodies for 1 hour at room temperature. Then ovaries were washed and mounted as above. Antibody dilutions were as follows: rabbit anti-Cid1 1:1000, rabbit anti-Cid5 1:1000, and Alexa Fluor goat anti-rabbit 568 1:1000.

### Embryo collection, fixation, immunofluorescence and imaging

To characterize Cid1 and Cid5 in the early embryo we imaged embryos produced from mothers and fathers with both Cid1GFP and Cid5mCherry transgenes. 0–60 min old embryos were collected on grape agar plates. Embryos were incubated in 30% bleach for 2 minutes to remove chorion. Fixation and antibody staining was performed according to Fanti and Pimpinelli method 3 (Fanti and Pimpinelli 2004). Briefly, embryos were transferred to a 1:1 mixture of heptane and methanol and shaken vigorously for one minute. The heptane layer was removed and embryos were washed twice with ice-cold methanol. Embryos rehydrated in PBS plus a drop of PBS + 0.1% Triton. Next, embryos were permeablized in PBS + 1% Triton for 30 min at room temperature. Embryos were blocked PBS + 0.1% TritonX + 3% BSA (BSA block) for one hour at room temperature. We diluted primary antibodies in BSA block and incubated overnight at 4°C. Embryos were washed and then incubated with secondary antibodies diluted in BSA block for two hours at room temperature. We washed embryos again after incubation with secondary antibodies and mounted embryos in wash solution (PBST). Embryos were imaged immediately after mounting. Primary antibody dilutions were the following: rabbit anti-AcH4 (Millipore, Billerica, MA; 1:1000), Alexa-Fluor goat secondary antibodies (Life Technologies) were diluted at 1:1000.

## Acknowledgements

We are grateful to Barbara Wakimoto for valuable discussions and advice. We would like to thank Ines Drinnenberg, Michelle Hays, Rini Kasinathan, Mia Levine, Antoine Molaro, Courtney Schroeder, and Janet Young for their comments on the manuscript. We thank the Drosophila Genetics Resource Center for the *D. virilis* tissue culture cells, Rainbow Transgenic Flies Inc. for the *D. virilis* transgenic injections, Covance Inc. for generating the polyclonal antibodies and the National Drosophila Species Stock Center (San Diego/ Cornell) for the *D. virilis* flies. This work was supported by funding from the National Institutes of Health training grants T32 HG000035 and T32 GM007270 (to L.E.K.) and R01 GM074108 (to H.S.M.). The funders played no role in study design, data collection and interpretation, or the decision to publish this study. H.S.M. is an Investigator of the Howard Hughes Medical Institute.

---

**Supplementary Figure 1. Cid1 and Cid5 antibody validation.** (A) Protein alignment of *D. virilis* Cid1 and Cid5. Peptides used to generate polyclonal antibodies are highlighted in green (Cid1) and magenta (Cid5). (B) Immunostaining of *D. virilis* larval brains with Cid1 (top) or Cid5 (bottom) antibodies. (C) *D. melanogaster* KC cells transfected with Venus-Cid5 overexpressed under the control of the *D. melanogaster* heat shock promoter. Venus and Cid5 antibody signal are shown. (D) A schematic indicating the site of GFP or mCherry insertion into Cid1 or Cid5, respectively. The bold ‘GGG’ represents a glycine linker that was added before and after each flurophore.

**Supplementary Figure 2. Localization of Cid1 and Cid5 in ovaries using antibodies.** Panels (A) – (C) and (G) use Cid1GFP and the Cid5 antibody to visualize Cid1 and Cid5 protein. Panels (D) – (F) and (H) use the Cid1 antibody and Cid5mCherry to visualize Cid1 and Cid5 protein. The schematics at the top of the figure indicate which protein detection methods were used. (A) A stage 8 egg chamber. Regions boxed in the right panel of (A) are shown at high magnification in subsequent panels. (B) High magnification image of a nurse cell from the stage 8 egg chamber boxed in (A). (C) High magnification image of the stage 8 oocyte boxed in (A). (D) A stage 8 egg chamber. (E) High magnification image of a nurse cell from the stage 8 egg chamber boxed in (D). (F) High magnification image of the stage 8 oocyte boxed in (D). (G) and (H) Image of a stage 14 oocyte nucleus in meiosis I metaphase arrest. Scale bars in (A) and (D) = 10μm. Scale bars in all other panels = 5μm.

**Supplementary Figure 3. Localization of Cid1GFP in testes.** Images showing Cid1GFP localization at various stages of spermatogenesis. (A) The apical tip (mitotic zone) of a *D. virilis* testis. Scale bar = 25μm (B) Two cells in late pro-metaphase or early metaphase. PH3S10 antibody staining is also shown. (C) Leaf-stage and (D) late-canoe stage sperm bundles. Scale bars = 5μm in (B) – (D). (E) Individualized mature sperm. Scale bar = 25μm

