## Supplementary figures and images for "Gametic specialization of centromeric histone paralogs in *Drosophila virilis*"

### Supplementary Figure 1

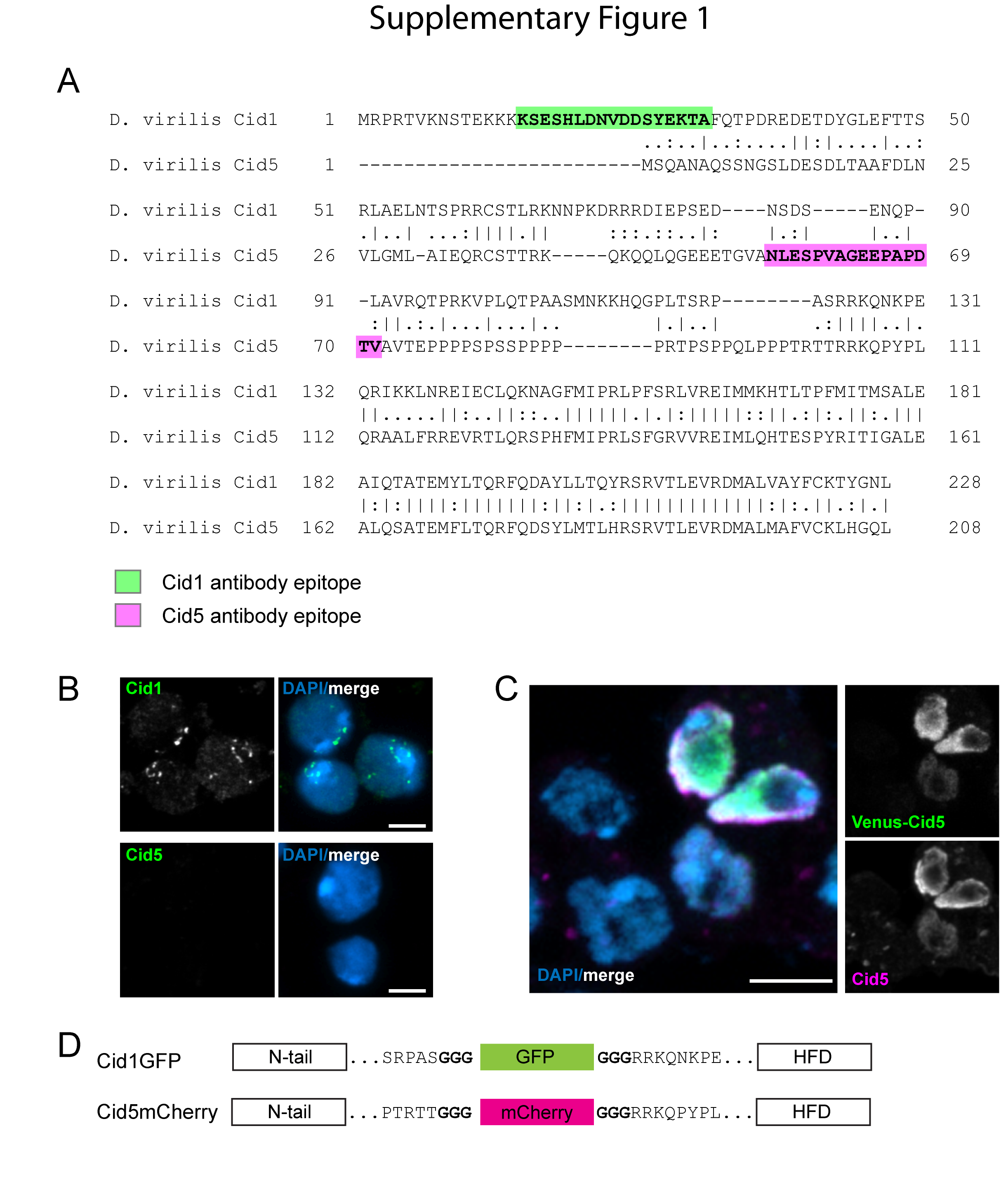

### Supplementary Figure 2

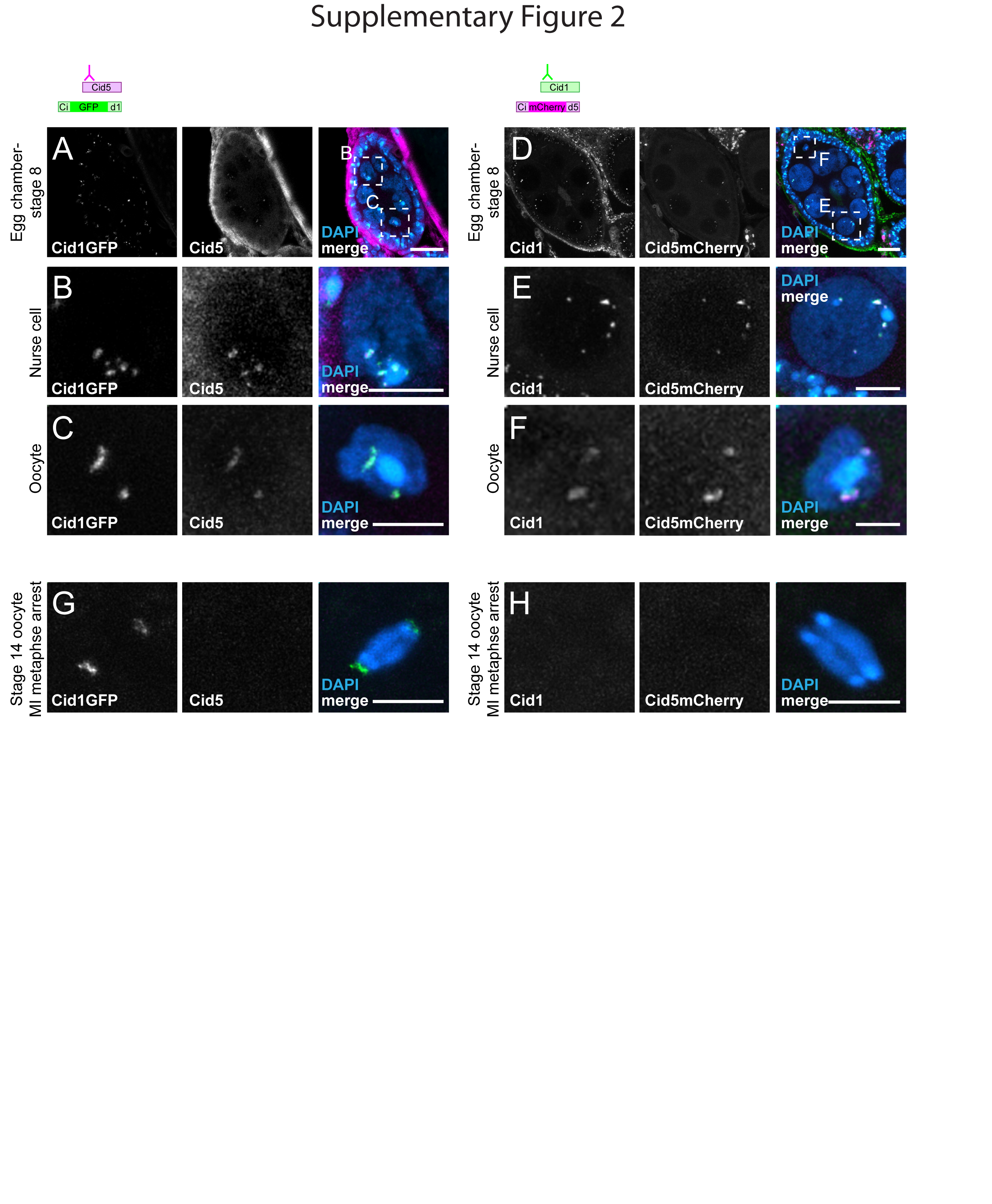

### Supplementary Figure 3

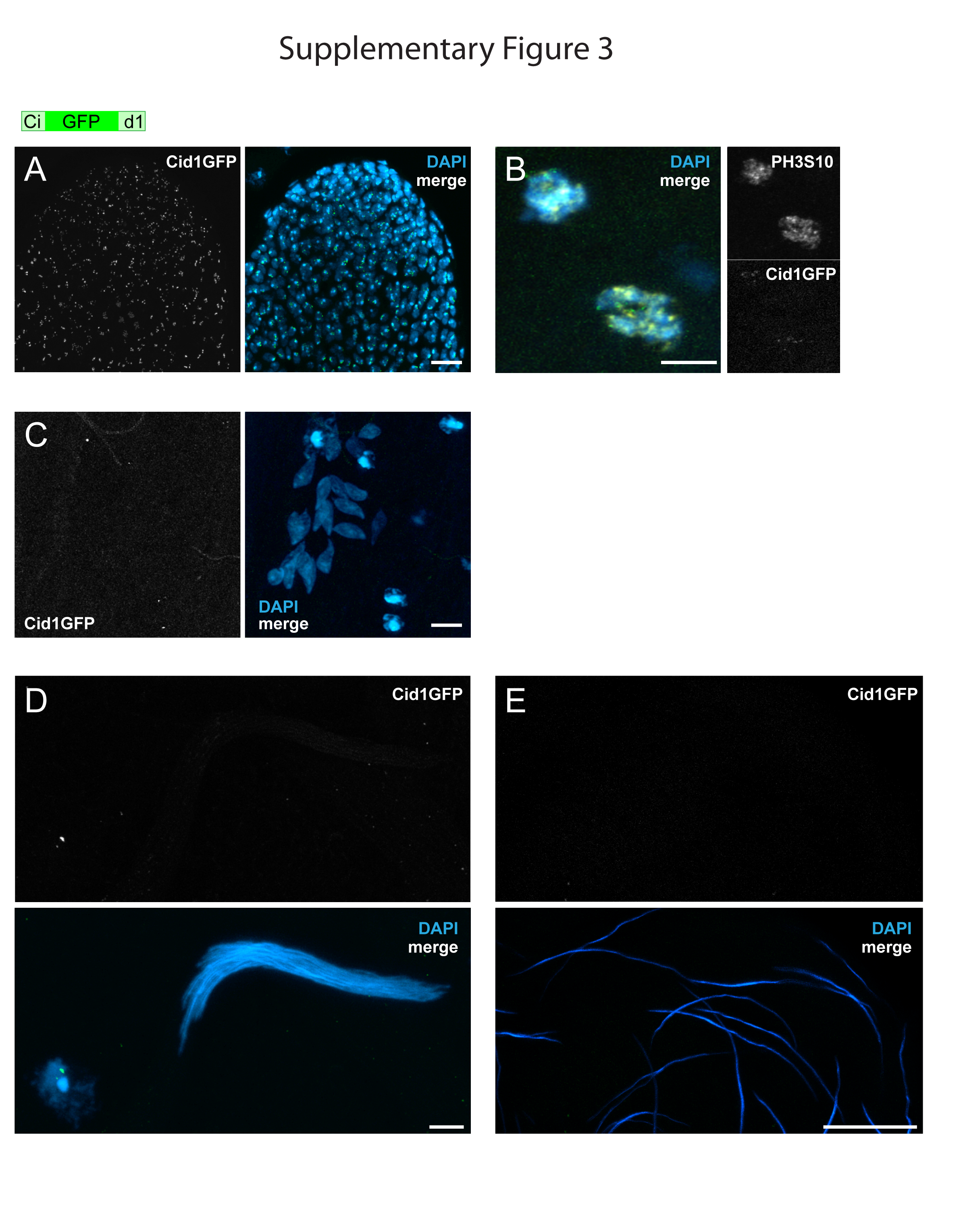
